# Human Induced Pluripotent Stem Cell Derived Sensory Neurons are Sensitive to the Neurotoxic Effects of Paclitaxel

**DOI:** 10.1101/2020.06.04.134262

**Authors:** Chenling Xiong, Katherina C. Chua, Tore B. Stage, Jeffrey Kim, Anne Altman-Merino, Daniel Chan, Krishna Saraf, Amanda Canato Ferracini, Faranak Fattahi, Deanna L. Kroetz

## Abstract

Chemotherapy-induced peripheral neuropathy (CIPN) is a dose-limiting adverse event associated with treatment with paclitaxel and other chemotherapeutic agents. The prevention and treatment of CIPN are limited by a lack of understanding of the molecular mechanisms underlying this toxicity. In the current study, a human induced pluripotent stem cell–derived sensory neuron (iPSC-SN) model was developed for the study of chemotherapy-induced neurotoxicity. The iPSC-SNs express proteins characteristic of nociceptor, mechanoreceptor and proprioceptor sensory neurons and show Ca^2+^ influx in response to capsaicin, α,β-meATP and glutamate. iPSC-SNs are relatively resistant to the cytotoxic effects of paclitaxel, with IC_50_ values of 38.1 μM (95% CI: 22.9 – 70.9 μM) for 48 hr exposure and 9.3 μM (95% CI: 5.7 – 16.5 μM) for 72 hr treatment. Paclitaxel causes dose- and time-dependent changes in neurite network complexity detected by βIII-tubulin staining and high content imaging. The IC_50_ for paclitaxel reduction of neurite area was 1.4 *μ*M (95% CI: 0.3 - 16.9 *μ*M) for 48 hr exposure and 0.6 *μ*M (95% CI: 0.09 - 9.9 *μ*M) for 72 hr exposure. Decreased mitochondrial membrane potential, slower movement of mitochondria down the neurites and changes in glutamate-induced neuronal excitability were also observed with paclitaxel exposure. The iPSC-SNs were also sensitive to docetaxel, vincristine and bortezomib. Collectively, these data support the use of iPSC-SNs for detailed mechanistic investigations of genes and pathways implicated in chemotherapy-induced neurotoxicity and the identification of novel therapeutic approaches for its prevention and treatment.

## Introduction

Chemotherapy induced peripheral neuropathy (CIPN) is a dose-limiting toxicity associated with a number of drugs used for the treatment of solid tumors and hematological cancers^1–3^. Drugs with diverse mechanisms of action, including microtubule disruptors, proteasome inhibitors and DNA-crosslinking agents all cause significant peripheral neuropathy. CIPN typically presents as burning, tingling or numbness in the hands and feet that occurs in a glove and stocking distribution^2, 4^. In addition to negatively affecting a patient’s quality of life, dose reductions, treatment delays and discontinuation can impact the therapeutic effectiveness of these drugs^2^. Despite years of research, there are no effective therapies to prevent and/or treat CIPN, highlighting the need to define the molecular basis of this toxicity to support the development of novel strategies for treatment.

The majority of the mechanistic studies of CIPN have employed behavioral testing in rodent models or cell-based studies using primary rodent dorsal root ganglion (DRG) neurons. Common mechanisms associated with the development of CIPN include axon degeneration, altered Ca^2+^ homeostasis, mitochondrial dysfunction, changes in neuronal excitability and neuroinflammation, although the relative contribution of these mechanisms varies for individual drugs^5–7^. For example, the microtubule stabilizing effects of paclitaxel inhibit anterograde and retrograde transport of synaptic vesicles down the microtubules, resulting in axon degeneration and membrane remodeling. This phenomenon is thought to be a major contributor to paclitaxel-induced peripheral neuropathy^8^. In contrast, the ability of DNA alkylators like cisplatin and oxaliplatin to form adducts with mitochondrial DNA and increase reactive oxygen species (ROS) contributes significantly to their peripheral neuropathy^5^. While these studies in preclinical models and primary cultures of rodent DRG neurons have enhanced our knowledge of potential mechanisms for CIPN, attempts to translate these findings into humans have been largely unsuccessful^3^.

In recent years, human induced pluripotent stem cell (iPSC)-derived neurons have been used for the study of CIPN. Commercially available iPSC-derived neurons (*e.g*., iCell Neurons^®^ and Peri.4U Neurons^®^) have been evaluated as a model of neurotoxicity^9^, used to screen for neurotoxic compounds^10–13^, and utilized for functional validation of genes identified in human genome wide association studies of CIPN^9, 14–16^. The use of human iPSC-derived neurons affords an advantage over rodent DRG neurons in their human origin and the potential to differentiate into specific peripheral sensory neuron populations. iCell^®^ neurons are a mixture of postmitotic GABAergic and glutamatergic cortical neurons that are more characteristic of relatively immature forebrain neurons than the sensory neurons found in the DRG^17, 18^. Peri.4U^®^ neurons are more peripheral-like, expressing βIII-tubulin, peripherin, MAP2 and vGLUT2, but have been minimally characterized with respect to functional properties^10, 19^. Additionally, neurons derived from human fibroblasts, blood and embryonic stem cells that express more canonical nociceptive markers like ISL1, BRN3A, P2RX3, the NTRK receptors and NF200^20–23^ have also been used to study chemotherapy toxicity. While these human derived cells resemble the DRG sensory neurons that are targeted by chemotherapeutics, there is significant interindividual variation across donor samples that limits their routine use for mechanistic studies and confounds the evaluation of functional consequences of genetic variation associated with human CIPN^24^.

Despite advances made in recent years in the development of human cell-based models for the study of CIPN, there remains a need for a robust, widely available and reproducible model for detailed mechanistic studies of this dose-limiting toxicity. The goal of the studies described below was to develop an iPSC-derived sensory neuron (iPSC-SN) model for the study of chemotherapy-induced neurotoxicity. Paclitaxel was used as a model neurotoxic chemotherapeutic to evaluate morphological, mitochondrial and functional changes associated with exposure of iPSC-SNs to neurotoxic compounds.

## Materials and Methods

### Neuronal differentiation

Human iPSC line WTC11^25^ was differentiated into sensory neurons following a published protocol^26^. An overview of the procedure is shown in Supplementary Figure S1 and detailed information about all reagents is found in Supplementary Table S1. Human iPSCs were originally cultured in mTesR medium (STEMCELL Technologies, Cambridge, MA) and more recently in StemFlex medium (ThermoFisher, Waltham, MA). iPSCs were cultured on matrigel (Corning, Corning, NY) coated plates at a seeding density of 50,000 cells/cm^2^. When iPSCs reached 80% confluency, neural differentiation (days 0-5) was initiated using KSR medium and SMAD inhibitors (100 nM LDN-193189 (Selleck, Houston, TX) and 10 μM SB431542 (Selleck, Houston, TX)). The KSR medium consisted of 80% knockout DMEM, 20% knockout serum replacement, 1X glutamax, 1X MEM nonessential amino acids and 0.01 mM β-mercaptoethanol (ThermoFisher, Waltham, MA). Sensory neuron differentiation began on day 2 with the addition of 3 μM CHIR99021, 10 μM SU5402, and 10 μM DAPT (Selleck, Houston, TX). N2 medium (ThermoFisher, Waltham, MA) was added in stepwise 25% increments every other day starting at day 4 (25% N2/75% KSR medium) through day 10 (100% N2 medium). N2 medium was composed of 50% DMEM/F12 medium with 100X N2 supplement (ThermoFisher, Waltham, MA) and 50% Neurobasal medium with 50X B27 supplement (ThermoFisher, Waltham, MA). On Day 12, differentiated sensory neurons were dissociated with Accutase (ThermoFisher, Waltham, MA) and replated onto 96-well plates (Greiner Bio-One, Monroe, NC), 6-well pates (Genesee Scientific, San Diego, CA), or μ-dishes or μ-slide chambers (ibidi, Fitchburg, WI). Neurons were plated at a density of 35,000 cells/cm^2^ in 96-well plates, μ-dishes and μ-slide chambers and 50,000 cells/cm^2^ for 6-well plates. All plates were triple coated with 15 μg/ml poly-L-ornithine hydrobromide (Sigma, St. Louis, MO), 2 μg/ml laminin (Fisher Scientific, Hanover Park, IL), and 2 μg/ml fibronectin (Fisher Scientific, Hanover Park, IL). Differentiated sensory neurons were maintained in N2 medium with neuronal growth factors (10 ng/ml human β-NGF, BDNF, NT3 and GDNF; PeproTech, Rocky Hill, NJ) in a 37°C incubator under 5% CO_2_. On day 15, cells were treated for 2 hr with freshly prepared mitomycin C (1 μg/ml) to eliminate non-neuronal cells. On day 17, medium was completely refreshed; subsequent 50% medium changes were made every 5-7 days. Sensory neurons were considered mature at day 35 and routinely used for experiments between days 35-45.

### Immunocytochemistry

Immunocytochemistry was performed on iPSC-SNs seeded in 96-well plates or μ-slide chambers. Cells were fixed with 4% paraformaldehyde in PBS for 15 min, permeabilized with PBS containing 0.1% Triton X-100 for 10 min, blocked with 5% goat serum in PBS for 1 hr and incubated overnight at 4°C with one of the following primary antibodies: rabbit anti-PAX6 (1:500, Covance), mouse/rabbit anti-Tubulin βIII clone TUJ1 (1:1000, Covance), goat anti-SOX10 (1:500, Santa Cruz), mouse anti-BRN3A (1:500, Millipore), goat anti-Peripherin (1:500, Santa Cruz), pig anti-TRPV1 (1:200, ThermoFisher) and rabbit anti-TrkA/B/C (1:50, Alomone Labs). Following several washes, the corresponding Alexa Fluor 488/568/594 - conjugated secondary antibodies (1:1000, ThermoFisher) were added for 1 hr at room temperature. Cells were incubated with DAPI (1:1000, ThermoFisher) for 10 min to visualize nuclei. A list of the primary and secondary antibodies used in this study is provided in Supplementary Table S2.

### Quantitative real-time PCR

Total RNA was isolated from iPSC-SNs seeded in 6-well plates using a RNeasy Mini Kit (Qiagen, Redwood City, CA) and reverse transcribed into cDNA using a SuperScript VILO cDNA Synthesis kit (Life Technologies, Grand Island, NY). Quantitative real-time PCR (qRT-PCR) was performed in 384-well reaction plates using 2X TaqMan Fast Universal Master Mix (Applied Biosystems, Foster City, CA), 20X TaqMan specific gene expression probes (Applied Biosystems, Foster City, CA; Supplementary Table S3), and 10 ng of the cDNA template. The reactions were carried out on an Applied Biosystems 7900HT Fast Real-Time PCR System (Applied Biosystems, Foster City, CA). The relative expression level of each mRNA transcript was calculated by the comparative ΔCt method^27^, normalized to the housekeeping gene hypoxanthine phosphoribosyl transferase (*HPRT*).

### Calcium imaging and analysis

Calcium imaging was performed on iPSC-SNs in 8-well μ-slide chambers. In some cases, sensory neurons were treated with 1 μM paclitaxel for 6-72 hr prior to Ca^2+^ imaging. Neurons were treated with 2 μM Fluo-4 AM (Life Technologies, Grand Island, NY) and 0.02% Pluronic F-127 (Life Technologies, Grand Island, NY) in HBSS for 15 min at 37°C. Cells were washed twice with warm HBSS before initiation of Ca^2+^ imaging using an inverted Nikon Ti microscope equipped with a CSU-W1 spinning disk confocal using a Plan Apo VC 20X/1.4 objective. Spontaneous Ca^2+^ transients were obtained at 37°C using a single-cell line scan mode with collection every 3 seconds. All imaging trials began with 60 s of baseline measurement before chemicals were added to the cells by micropipette. In a single experiment, neurons were exposed sequentially to channel and receptor agonists followed by 35 mM KCl. Agonists included capsaicin (10 μM), α,β-meATP (50 μM) and glutamate (100 μM). Analysis of calcium imaging videos was performed using ImageJ (NIH, Bethesda, MD). Regions of interest were drawn around at least 80 randomly selected soma. Mean pixel intensities were measured across the entire time-lapse. Each value was normalized to baseline on a cell-by-cell level and expressed as ΔF/F_0_ using the following equation:

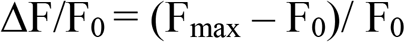

where F_max_ is the maximum intensity extracted for each cell following individual stimuli and F_0_ is the average intensity collected during the 60 s baseline period before addition of any stimuli.

### Sensory neuron treatment

Paclitaxel, vincristine, docetaxel, and hydroxyurea were purchased from Sigma-Aldrich (St. Louis, MO). Bortezomib was purchased from Selleck Chemicals (Houston, TX). Stock solutions were prepared in 100% dimethylsulfoxide (DMSO). Final drug concentrations for each experiment are indicated in the figures. The concentration of DMSO was maintained at 0.2% in all experiments.

### Viability and apoptosis assays

Cell viability was determined by measuring ATP levels and apoptosis by caspase 3/7 activation following paclitaxel treatment (0.001 – 50 μM) of mature iPSC-SNs for 24, 48 or 72 hr. ATP levels were assessed using the CellTiter-Glo Luminescent Cell Viability Assay (Promega, Madison, WI) according to the manufacturer’s instructions. Apoptosis was determined for the same paclitaxel treatments using the Caspase-Glo 3/7 assay (Promega, Madison, WI). Luminescence was measured on a Synergy 2 microplate reader (BioTek, Winooski, VT). At least six wells per drug dose for both assays were performed; cytotoxicity assays were repeated in cells from three separate differentiations and the apoptosis assay from two differentiations.

### Imaging and neurite analysis

Following immunostaining of the sensory neurons, the 96-well plates were scanned on an IN Cell Analyzer 6000 (GE Healthcare Life Sciences, Amersham, United Kingdom). A 10X objective provided sufficient resolution to distinguish neurite networks. Nine individual fields per well with a field-of-view of 15.9 mm^2^ (47% of well) were imaged in two channels. Images were batch processed through an imaging processing software, MIPAR^®^ (Worthington, OH), with a custom-built algorithm to analyze measurements for chemotherapy-induced neuronal damage (Supplementary Figure S2). The algorithm generates optimized grayscale images by reducing overall noise and minimizing the amount of non-specific staining to identify and quantify the neurite networks within each field-of-view image. A subsequent segmentation algorithm was performed to identify and quantify nuclei within each field-of-view image. After processing, each image yielded measurements of total neurite area and neuron count. Neurite area was defined by the total area of pixels captured within the βIII tubulin-stained network. Nuclei were used as the measure of neuron number, rejecting DAPI-stained particles less than 50 pixels to exclude non-specific staining. To get a global measurement for each well, total neurite area and total cell count were generated by summing measurements across the nine field-of-view images. Per-well images were stitched together using an in-house script to batch process all nine field-of-view images, using the Grid/Collection Stitching plugin in Fiji. Processed images included further in the analysis were required to pass quality control on a per-well basis to assess the quality of the neurons and images. Wells were only included if 1) neurites covered ≥ 50% of the entire well, 2) no more than 3 field-of-view images (out of 9) were out-of-focus, and 3) a majority of the signal intensities captured were not from artifacts. Per-well quality control was performed manually by at least two investigators blinded to the treatments. For each experiment, all drug treatments had 6-8 replicates and raw neurite area measurements and cell counts from imaging data were averaged to obtain a mean total neurite area and cell count for each condition. Experiments were repeated at least three times and the reported values represent mean phenotype measurements from independent neuron differentiations.

### Mitochondrial membrane potential and mobility

Mature sensory neurons in 8-well μ-slide chambers were co-stained with 10 nM tetramethylrhodamine methyl ester (TMRM; Life Technologies, Grand Island, NY) and 100 nM MitoTracker Green FM (Life Technologies, Grand Island, NY) for 30 min. Fluorescence imaging was performed using an inverted Nikon Ti microscope equipped with a CSU-W1 spinning disk confocal using a Plan Apo VC 100X/1.4 water objective with incubation system. The cells were kept at 37°C in 5% CO_2_ during imaging. MitoTracker Green FM was used to localize the mitochondria. Mitochondrial membrane potential (ΔΨ_m_) was quantified as the TMRM intensity normalized to MitoTracker Green FM intensity. Mitochondrial mobility in neurites was recorded by time-lapse imaging; images were taken every 3 sec for 6 min with a 100 ms exposure time. Mitochondrial fluorescent signals were quantified with ImageJ. Measurements were repeated in neurons from three different differentiations.

### Statistical Analysis

Mean neurite areas and mean cell counts for each experiment were expressed as a ratio of drug-treated to vehicle-treated samples and were compared across at least three differentiations. Normality of relative ratios was confirmed using a Shapiro-Wilk test. Differences between relative ratios for the treatment groups were tested for significance by ordinary one-way ANOVA and subsequent Dunnett post hoc comparisons to controls using Prism software (GraphPad, La Jolla, CA). Calcium and mitochondria flourescence images were quantified by ImageJ and analyzed by parametric one-way ANOVA and Dunnett post hoc comparisons to controls. Statistical significance was set at *p* < 0.05.

## Results

### iPSC differentiation into sensory neurons

Human iPSCs were differentiated into a sensory neuron lineage following a published protocol^26^. The cells showed expression of the neuroectoderm marker PAX6, the neural crest marker SOX10, the neuronal marker TUBB3 and the canonical sensory neuron markers BRN3A, peripherin and TRPV1 at the expected times during differentiation (Figure 1a). Phase contrast images in mature sensory neurons demonstrate ganglia-like structures and typical neurite extensions (Figure 1a). RNA levels of each marker gene throughout differentiation and maturation were consistent with the immunostaining (Figure 1b). There was a rapid loss of the pluripotency factors NANOG and OCT4 upon initiation of differentiation that coincided with an increase in the expression of PAX6, a neuroectoderm marker. The neural crest marker SOX10 peaked at day 12 and decreased after day 16. TUBB3 expression peaked at day 8 and remained elevated through day 50. The nociceptor neuron markers TAC1 and NTRK1 were also detected after a week of differentiation and remained elevated. The fully differentiated peripheral sensory neurons at day 35 expressed all three NGF receptor family members (TRKA, TRKB and TRKC encoded by *NTRK1*, *NTRK2* and *NTRK3*, respectively) (Figure 1c), indicating that the sensory neurons were likely comprised of mechanoreceptor, nociceptor and proprioceptor subtypes.

**Figure 1.**
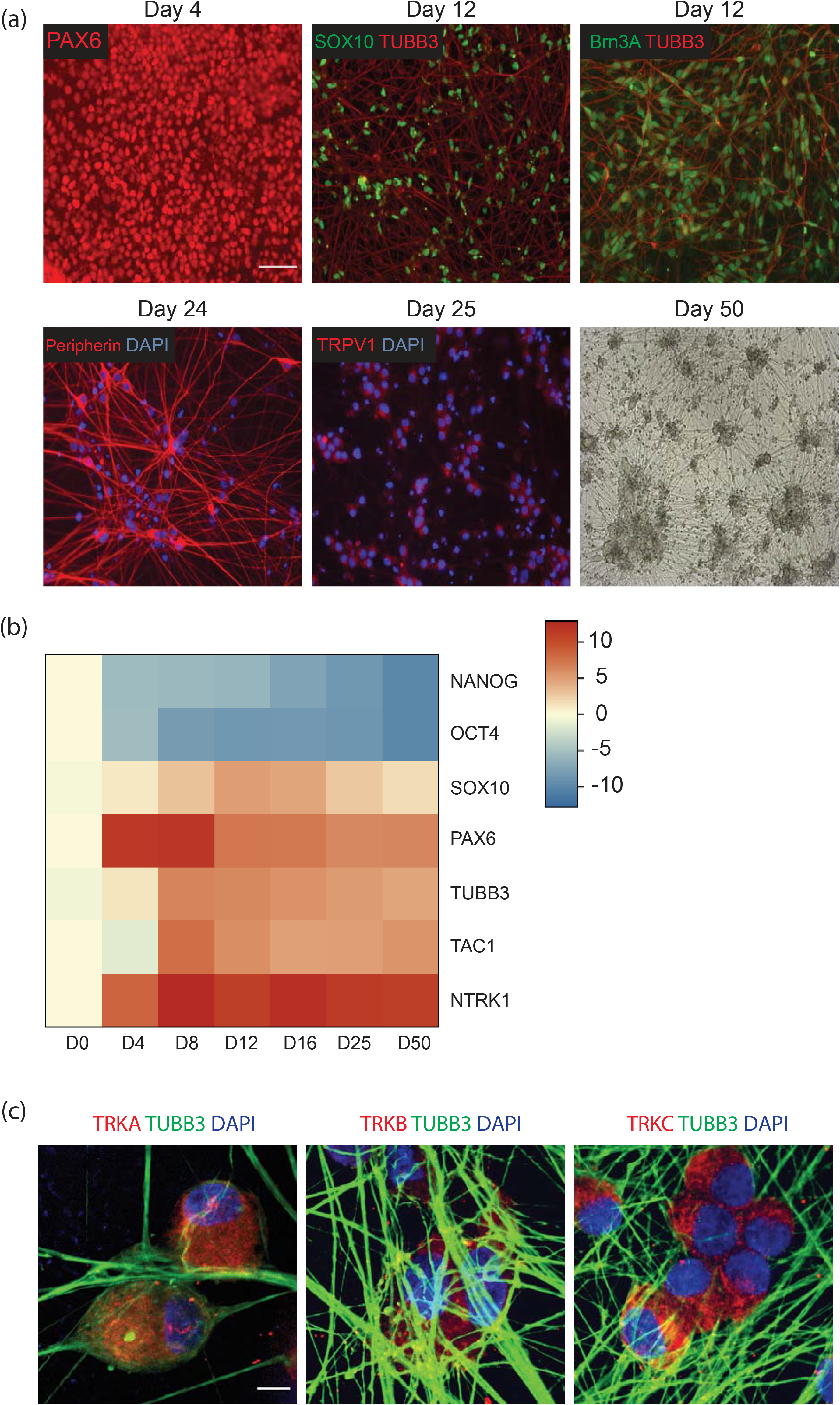
Differentiation of human iPSCs into sensory neurons. iPSCs were differentiated using a cocktail of small molecule inhibitors. (a) At the indicated times, cells were fixed and stained with the neuroectoderm marker PAX6, the neural crest marker SOX10, the neuron marker TUBB3, and the sensory neuron markers Brn3A, peripherin and TRPV1. A phase contrast image shows the characteristic ganglia and neurite networks. Scale bar, 100 μm (b) Heat map of qRT-PCR results showing the relative expression levels of key marker genes during differentiation on a log_2_ scale. Each column represents the mean expression levels of three independent samples at the indicated time points. (c) Immunocytochemistry of Day 35 peripheral sensory neurons for the subtype specific markers TRKA, TRKB and TRKC; scale bar, 10 μm.

Calcium imaging was performed to investigate the functional properties of the induced sensory neurons using the fluorescent marker Fluo-4 AM. Only cells with a stable baseline and a characteristic increase in Ca^2+^ flux in response to KCl were evaluated. Agonists for TRPV1 (10 μM capsaicin), P2X3 (50 μM α,β-meATP), and glutamate receptors (100 μM glutamate) were tested for selective activation of their respective cognate receptors. Representative images from treatments are shown in Figure 2a. Both capsaicin and α,β-meATP evoked calcium transients in subpopulations of iPSC-SNs (Figures 2b and 2c), demonstrating functional activity of nociceptive sensory neurons. A large number of mature sensory neurons also showed high calcium flux in response to glutamate (Figures 2b and 2c), indicating an excitatory glutamatergic neuronal phenotype.

**Figure 2.**
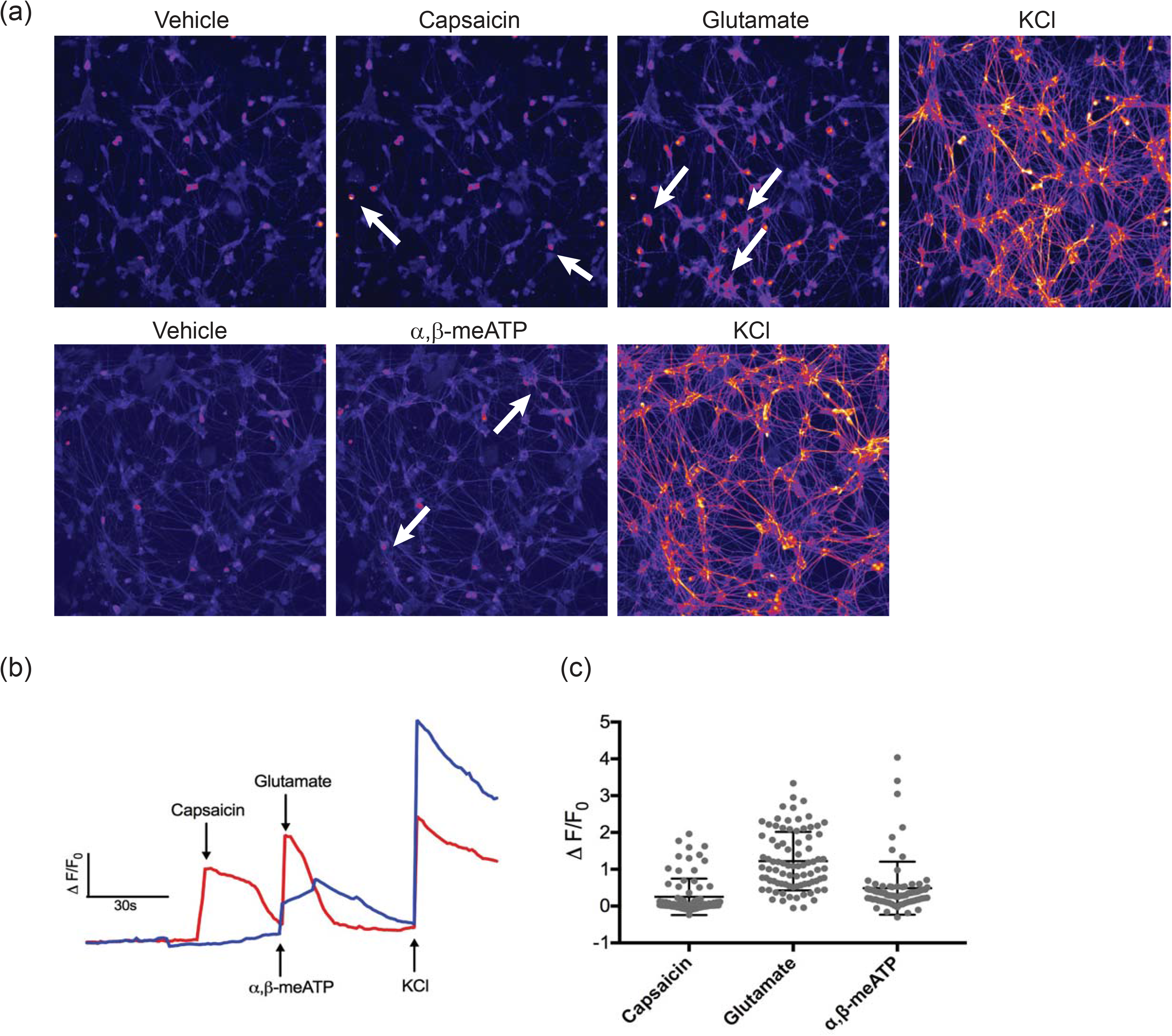
Response of iPSC-SNs to agonists of sensory neuron channels and receptors. iPSC-SNs were treated with vehicle (0.2% DMSO) or selective channel/receptor agonists. (a) Representative images showing Ca^2+^ signals (red) after treatment with 10 *μ*M capsaicin, 100 *μ*M glutamate, 50 *μ*M α,β-meATP or 35 mM KCl. Arrows indicate examples of calcium flux induced by individual stimuli. Scale bar, 100 μm (b) Representative fluorescence trace of calcium flux in a single cell. Values are plotted as the change in fluorescence intensity from baseline divided by the baseline (ΔF/F_0_). Data for each condition were collected sequentially at the indicated time points: capsaicin (45 s), glutamate and α,β-meATP (72 s), and KCl (120 s). (c) Scatter dot plots of ΔF/F_0_ of individual cells after application of agonists. Individual dots represent the response of functional neurons (responsive to KCl) to capsaicin, glutamate or α,β-meATP. At least 80 functional neurons were analyzed across three independent differentiations.

### Paclitaxel cytotoxicity

Investigation of the molecular mechanisms underlying paclitaxel-induced neurotoxicity in the iPSC-SNs requires conditions where neuron morphology and function are disrupted without overt cytotoxicity at clinically relevant concentrations (≤10 μM)^28^. Treatment with up to 10 *μ*M paclitaxel for 24 hr had no significant effect on cell viability (Figure 3a). There was a 20-30% loss in cellular ATP with exposure to 0.5 *μ*M paclitaxel for 48 or 72 hr, and ATP levels gradually decrease with concentrations of paclitaxel ≥5 μM. Treatment with 50 μM paclitaxel for 48 or 72 hr showed significant effects on cell viability, with >50% decrease in cellular ATP levels. The IC_50_ values for paclitaxel cytotoxicity at 48 hr and 72 hr are 38.1 μM (95% CI: 22.9 – 70.9 μM) and 9.3 μM (95% CI: 5.7 – 16.5 μM), respectively. Similar to cytotoxicity, exposure of the iPSC-SNs to paclitaxel concentrations as high as 50 μM for 24 hr did not induce apoptosis (Figure 3b). Exposure of the iPSC-SNs with up to 1 μM paclitaxel for 48 or 72 hr increased caspase 3/7 activity less than 25% compared to the vehicle control but treatment with higher concentrations for 48 or 72 hr increased apoptosis (Figure 3b).

**Figure 3.**
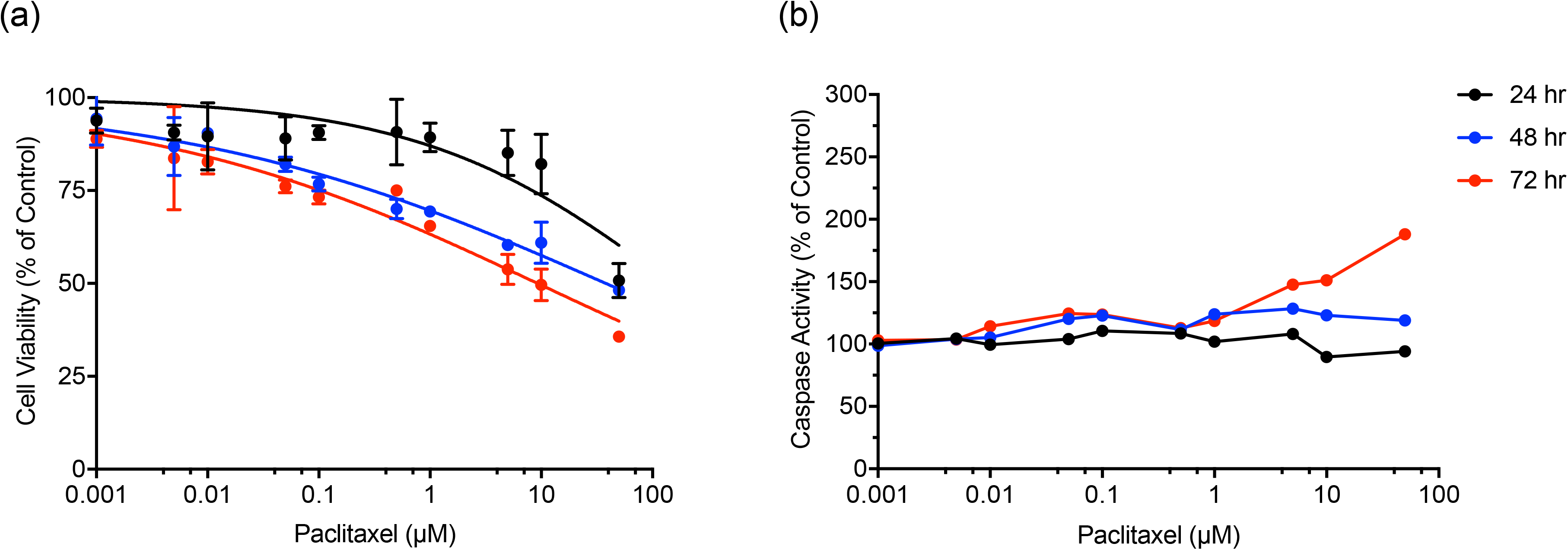
Effect of paclitaxel on cell viability and apoptosis. iPSC-SNs were treated for 24, 48 or 72 hr with the indicated concentrations of paclitaxel. Cellular levels of ATP were measured as an indicator of cell viability (a) and caspase 3/7 activity was measured to detect apoptosis (b). ATP levels and caspase 3/7 activity are expressed relative to DMSO-treated cells. The values shown for cell viability are the mean ± SD from three differentiations with the line representing the dose-response data fit to a Hill equation. The mean value for two differentiations are shown for the apoptosis assay; the percent difference in normalized apoptosis values between the two differentiations averaged 15-20%. At least six wells per drug dose were used for both assays with cells from a single differentiation.

### Concentration- and time-dependent effects of paclitaxel on neurite morphology

Because chemotherapy-induced neuropathy is characterized by a distinctive axon degeneration, neurite morphology changes were assessed as a marker of neurotoxicity. Mature iPSC-SNs had complex neurite networks extending from their cell bodies that were visible by staining of microtubules with a neuron specific βIII-tubulin antibody (Figure 4a). Both time- and concentration-dependent changes in the complexity of these networks were observed following treatment of iPSC-SNs with paclitaxel (Figure 4a). Hydroxyurea was used as a negative control since it is a cytotoxic chemotherapy drug without known neurotoxicity, and treatment with 1 μM hydroxyurea had no visible effect on neurite networks. Vincristine, a known neurotoxic agent, was included as a positive control; neurite networks were completely disrupted with 48-72 hr exposure to 0.1 μM vincristine. Total neurite staining for βIII-tubulin was quantified as a measure of neurite complexity and reported as neurite area. Treatment with 0.001-10 *μ*M paclitaxel for 24 hr had no significant effect on neurite area compared to controls (Figure 4b), while treatment with 1 μM and 10 μM paclitaxel for 48 hr reduced neurite area relative to vehicle control by 45% (*p* = 0.001) and 69% (*p* = 0.007), respectively (Figure 4b). Longer exposure (72 hr) of iPSC-SNs to 1 μM and 10 μM paclitaxel reduced neurite area by 57% (*p* = 0.0365) and 73% (*p* = 0.0083), respectively (Figure 4b). The IC_50_ for paclitaxel reduction of neurite complexity was 1.4 *μ*M (95% CI: 0.3 - 16.9 *μ*M) for 48 hr exposure and 0.6 *μ*M (95% CI: 0.09 - 9.9 *μ*M) for 72 hr exposure. There was no significant effect of paclitaxel treatment on number of neurons (Figure 4c; *p* > 0.05), consistent with the limited cytotoxicity and apoptosis at these concentrations (Figure 3). The negative control hydroxyurea had no effect on neurite complexity or number of neurons (Figures 4b and 4c), while 0.1 μM vincristine significantly reduced neurite area (Figure 4b). Variability in neuron numbers with vincristine treatment (Figure 4c) is likely an artifact of the imaging analysis under conditions of significant cytotoxicity.

**Figure 4.**
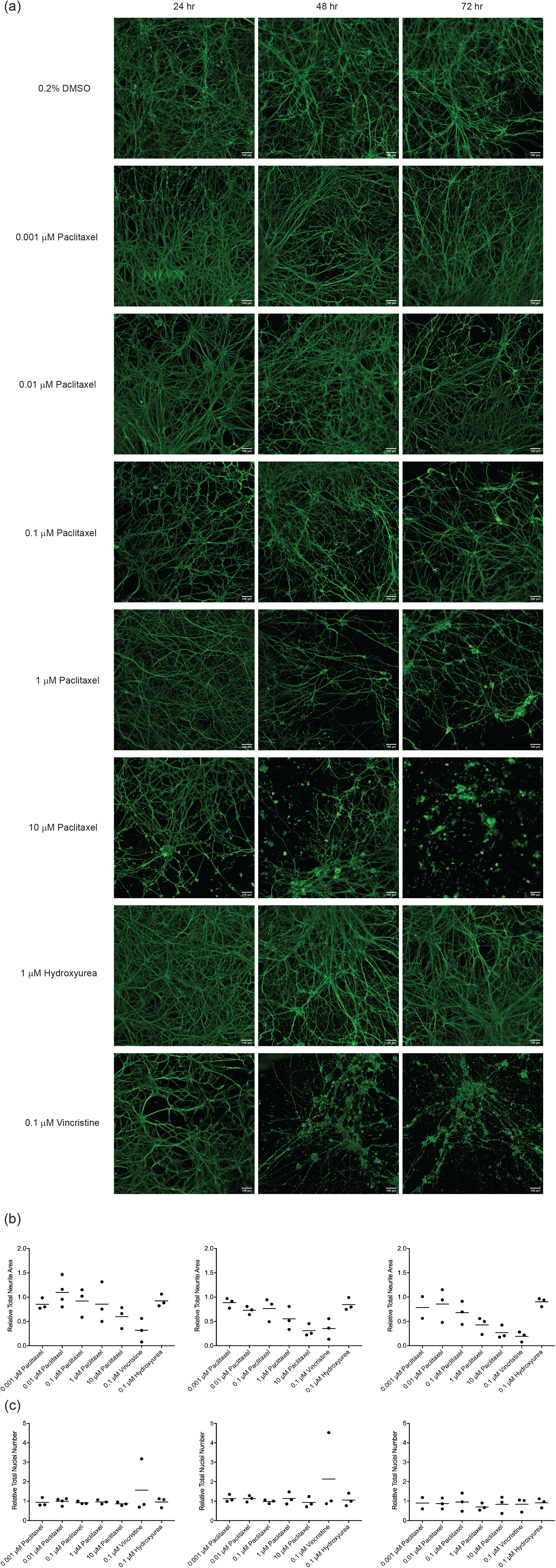
Concentration- and time-dependent effects of paclitaxel on iPSC-SN neurite networks. Mature iPSC-SNs were treated with the indicated concentrations of paclitaxel for 24, 48 or 72 hr. Controls were treated with vehicle (0.2% DMSO), a non-neurotoxic drug (hydroxyurea) or a potent neurotoxin (vincristine). (a) Representative images are shown for the indicated treatments. Cells were fixed and stained for neuron specific βIII-tubulin (green). Images were taken at 10X magnification, scale bar 100 μm. (b) Neurite area (βIII-tubulin staining) and (c) neuron number (DAPI staining) were quantified following drug exposure using high content image analysis and are expressed relative to the DMSO control. Each data point represents the mean measurement of 6-8 replicates from a single independent differentiation expressed relative to vehicle controls. The coefficient of variation in vehicle-treated neurites ranged from 10-28%. Relative mean neurite area and neuron counts at each exposure time were tested for differences across drug conditions by one-way ANOVA with Dunnett post hoc comparisons to controls. Paclitaxel treatment with 1 and 10 μM paclitaxel for 48 and 72 hr caused significant loss of neurite area (*p* < 0.05).

### Paclitaxel-induced changes in Ca^2+^ flux

Mature sensory neurons (day 35) were treated with 1 μM paclitaxel for 6, 24 or 72 hr to assess changes in neuron excitability. Live Ca^2+^ imaging was performed both at baseline and upon sequential treatment with 100 μM glutamate and 35 mM KCl. iPSC-SNs treated with paclitaxel displayed significant changes in Ca^2+^ flux in response to glutamate-induced depolarization compared to vehicle treated cells (Figure 5; *p* < 0.0001). With short-term exposure to paclitaxel (6 hr), the mean fluorescence intensity increased 44% in response to glutamate (95% CI: 24-63%, *p* = 0.0001, Dunnet’s test). Exposure to paclitaxel for 24 hr caused a 58% decrease in glutamate-induced Ca^2+^ flux (95% CI: 47-68%, *p* = 0.0001, Dunnet’s test) and 72 hr exposure decreased glutamate-induced Ca^2+^ flux 77% (95% CI: 66-88%, *p* = 0.0001, Dunnet’s test).

**Figure 5.**
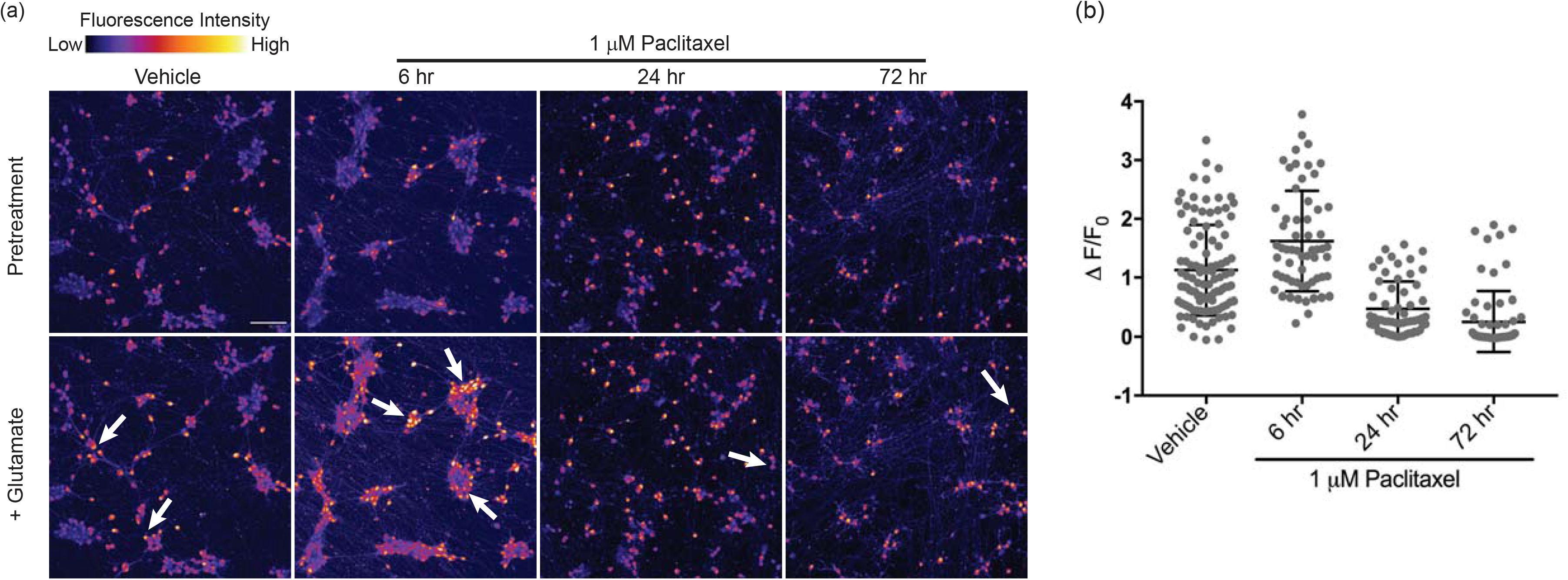
Response of paclitaxel treated iPSC-SNs to glutamate. Mature iPSC-SNs were pretreated with 0.2% DMSO, or 1 μM paclitaxel for 6, 24 or 72 hr. Calcium flux images were taken prior to and after addition of 100 μM glutamate. (a) Representative images of calcium flux before and after glutamate treatment are shown. Scale bar, 100 μm (b) ΔF/F_0_ intensity plot showing the response of individual cells after application of glutamate. Depolarization with 35 mM KCl was used at the end of each experiment to confirm neural identity and sustained functional capacity.

### Paclitaxel-induced changes in mitochondrial membrane potential

Mitochondrial dysfunction and oxidative stress have been implicated in the pathophysiology of CIPN^29^. To determine whether paclitaxel induced mitochondrial changes in iPSC-SNs, ΔΨ_m_ was measured using the ΔΨ_m_-dependent probe TMRM. The ΔΨ_m_-independent probe Mitotracker Green FM was used to visualize and analyze mitochondrial distribution and movement along the neurites (Figure 6a). Mitochondria were widely distributed along the neurites and cell bodies, with no apparent difference in distribution between the cells treated for 72 hr with vehicle or paclitaxel (Figure 6a). Mitochondrial membrane potential normalized to mitochondrial density decreased in iPSC-SNs treated with paclitaxel (*p* = 0.0368, ANOVA). Following treatment with 1 μM paclitaxel for 72 hr, normalized ΔΨ_m_ decreased 41% (95% CI: 1-82%) compared to vehicle treated controls (*p* = 0.0417, Dunnett’s test) (Figure 6b). Mitochondrial movement in both directions was also reduced in paclitaxel-treated iPSC-SNs, with almost complete suppression of mitochondrial trafficking down the neurites with exposure to 1 μM paclitaxel for 72 hr (Supplementary Movies).

**Figure 6.**
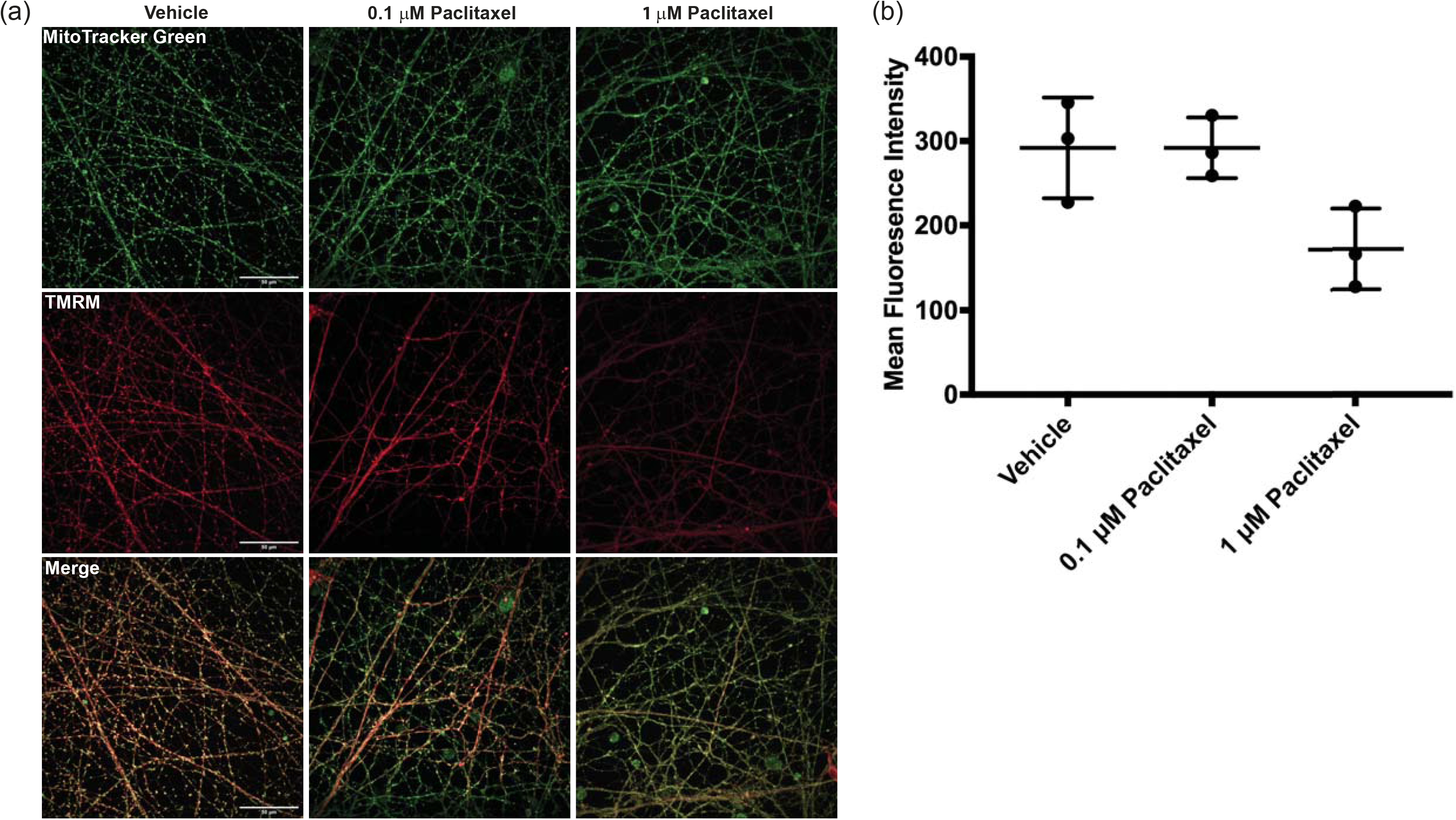
Alterations in mitochondrial membrane potential after paclitaxel treatment. Mature iPSC-SNs were treated with vehicle or paclitaxel (0.1 or 1 μM) for 72 hr. (a) Cells were stained with MitoTracker Green FM to detect mitochondria and with TMRM to measure mitochondrial membrane potential. Representative images from three independent differentiations are shown. Scale bar, 50 μm (b) Mean fluorescence intensity of the mitochondrial membrane potential was quantified by ImageJ and is shown as individual values for each experiment, with the mean and SD. The intensities were tested across conditions by one-way ANOVA with post-hoc comparisons. Compared to the vehicle, fluorescence intensity is significantly decreased upon 1 μM paclitaxel treatment for 72 hr (*p* = 0.0417).

### Differential sensitivity of iPSC-SNs to chemotherapeutic drugs

Since multiple chemotherapeutic drugs cause sensory peripheral neuropathy, the effect of additional agents on neurite complexity was assessed. Concentrations of each drug needed for robust quantification of changes in neurite area were selected based on published data^13, 19^ and treatment was restricted to 72 hr based on the data from paclitaxel presented above. As illustrated in Figure 7, vincristine caused the largest decrease in neurite area relative to vehicle control, with a 54% and 75% (*p* = 0.0001) decrease following exposure to 0.01 μM and 0.1 μM vincristine, respectively. Bortezomib decreased neurite area to a similar degree at concentrations of 0.1 and 1 μM (46% (*p=* 0.0004) and 51% (*p* = 0.0001) decrease, respectively). Docetaxel and paclitaxel had similar neurotoxicity, with a 40-45% decrease in neurite area (*p=* 0.0018, docetaxel; *p* = 0.0004, paclitaxel) following exposure to 1 μM and 27-30% decrease (*p*= 0.0446, docetaxel; *p* = 0.0237, paclitaxel) at 0.1 μM (Figure 7b). Hydroxyurea had no effect on neurite complexity at either concentration tested (*p* > 0.05). None of the drug treatments had an effect on cell number (*p* > 0.05), indicating that the changes in neurite area were not due to cytotoxicity (Figure 7c).

**Figure 7.**
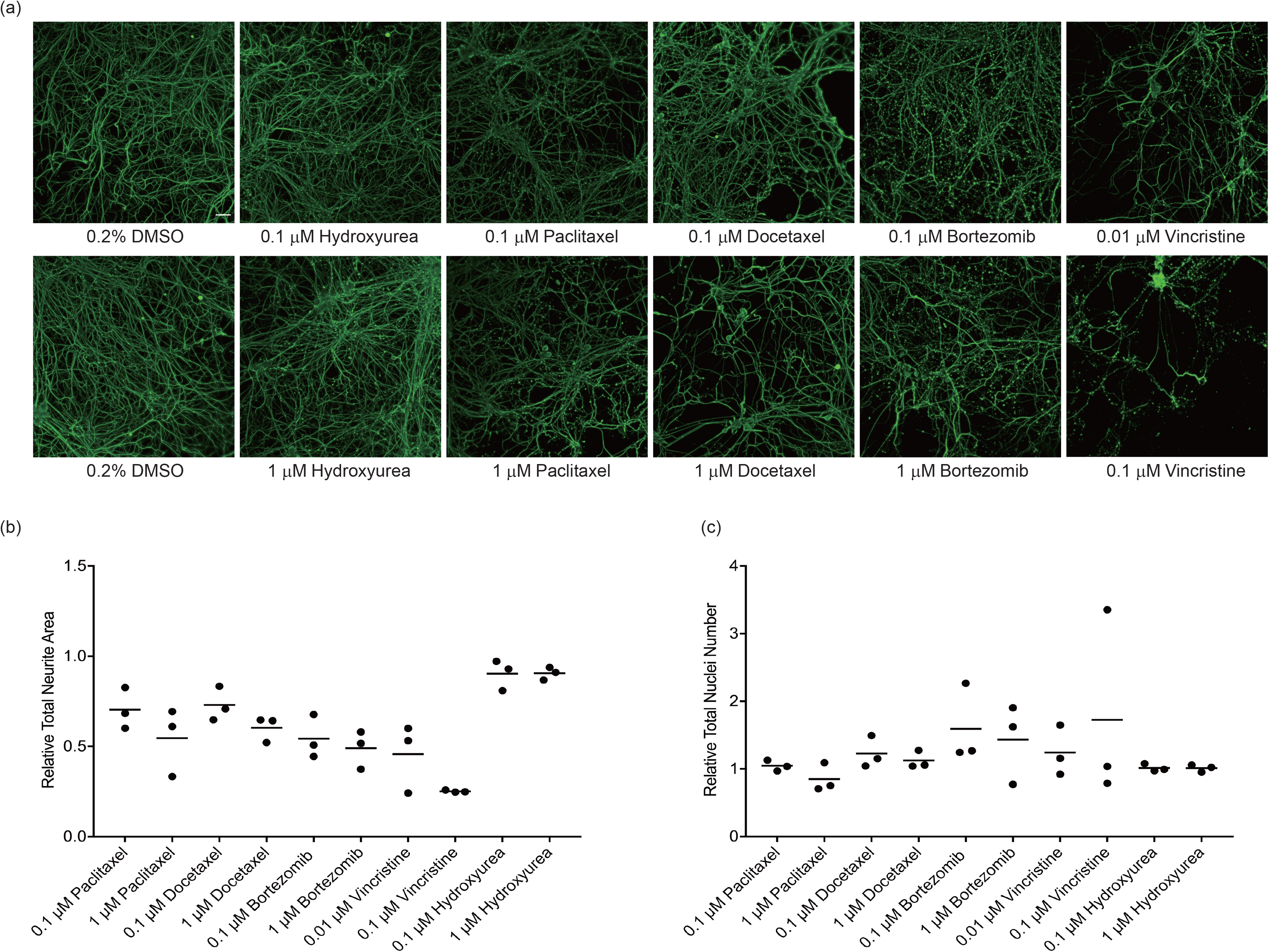
Effect of chemotherapy agents on neurite morphology in iPSC-SNs. Mature iPSC-SNs were treated with the indicated concentrations of paclitaxel, docetaxel, bortezomib, vincristine or hydroxyurea for 72 hr; controls were treated with vehicle (0.2% DMSO). (a) Representative images are shown for the indicated treatments. Cells were fixed and stained for neuron specific βIII-tubulin (green). Images were taken at 10X magnification, scale bar 100 μm. (b) Neurite area (βIII-tubulin staining) and (c) neuron number (DAPI staining) were quantified following drug exposure using high content image analysis and are expressed relative to the DMSO control. Each data point represents the mean measurement of 6-8 replicates from a single independent differentiation expressed relative to vehicle controls. The coefficient of variation in vehicle-treated neurites ranged from 11-24%. Relative mean neurite area and neuron counts at each exposure time were tested for differences across drug conditions by one-way ANOVA with Dunnett post hoc comparisons to controls. All treatments except hydroxyurea caused a significant loss of neurite area relative to vehicle-treated cells (*p* < 0.05).

## Discussion

A model of human sensory neurons derived from induced pluripotent stem cells was developed to investigate the mechanistic basis of chemotherapy-induced peripheral neuropathy. Mature iPSC-SNs display ganglia-like structures and complex neurite networks, express canonical markers of peripheral sensory neurons, and respond to agonists of critical channels and receptors. Importantly, paclitaxel-induced changes in neurite networks, mitochondrial function and glutamate receptor signaling were consistent with studies of this chemotherapeutic in preclinical models and rodent DRG neurons^5–7^. The iPSC-SNs are sensitive to a diverse group of neurotoxic chemotherapeutics and will be a valuable model to identify genes and pathways contributing to dose-limiting CIPN.

The derivation of human sensory neurons from embryonic stem cells (ESCs) and iPSCs was initially described using a combination of dual SMAD inhibition to generate neural crest cells and Wnt activation coupled with inhibition of Notch, VEGF, FGF and PDGF signalingto generate sensory neurons^26^. The sensory neurons were subsequently induced to mature nociceptors by culturing with neurotrophic factors. The iPSCs in the current study were similarly differentiated through a SOX10 positive neural crest intermediate and expressed peripheral neuron markers including TUBB3, BRN3A and TAC1 by day 10-12 of differentiation. Other similarities include the ganglia-like clusters that develop with extensive neurite formation and the response of subsets of iPSC-SNs to the TRPV1 activator capsaicin and the P2RX3 activator α,β-meATP. The current iPSC-SNs do show some differences to those described by Chambers et al. ^26^; most notable is the expression of not only TRKA (NTRK1), but also TRKB (NTRK2) and TRKC (NTRK3) subtypes. This suggests that the iPSC-SNs described in this report are a mixture of TRKA^+^ nociceptor, TRKB^+^ mechanoreceptor, and TRKC^+^ proprioceptor sensory neurons.

Heterogeneity in sensory neuron subtypes derived from human ESCs and iPSCs has been previously reported. In an extensive characterization of 107 iPSC lines differentiated and induced into sensory neurons using the same protocol as the current study, the expression of NTRK1-3 were detected by RNA-seq^24^. In contrast, neuroectodermal spheres from human ESCs and iPSCs were differentiated into neural crest cells and induced into sensory neurons that expressed almost exclusively NTRK2 and yielded functional low threshold mechanoreceptors^30^. A BRN3A and NGN2 transcription factor-driven differentiation of iPSCs was recently described and yields a homogeneous population of sensory neurons that express mostly NTRK1 and co-express TRPM8 and PIEZO2; these sensory neurons function as low threshold mechanoreceptors that also respond to cold stimuli^31^. Interestingly, application of the BRN3A/NGN2 transcription factor differentiation to a neural crest population derived from neuroectodermal spheres yielded two distinct sensory neuron populations; one expresses TRPM8 and PIEZO2 sensory neurons and a separate population derived by only transient BRN3A/NGN2 expression followed by culturing with neurotrophic factors expresses high levels of PIEZO2 but not TRPM8^31^. While there are clear differences in sensory neuron populations related to the method of differentiation and induction^24, 26, 30–32^, significant variability in iPSC-derived models of human disease has also been attributed to the iPSC genetic background, somatic mutations and variations in their culturing and maintenance^33^. The largest source of variation among sensory neurons derived from a large number of iPSC lines in a single institution with a standardized protocol was batch-to-batch variation across differentiations^24^. Careful attention to changes in cellular behavior during differentiation and induction are necessary to maintain reproducible conditions across differentiations. Despite these potential batch-to-batch differences, the results from the current study were collected over several years and the neurotoxicity phenotypes of interest were reproducible over this period. Future studies will address the effect of paclitaxel and other chemotherapeutic agents in more homogeneous sensory neuron populations, such as the cold and mechanosensitive neurons described recently^31^.

The iPSC-SNs behaved functionally as peripheral sensory neurons. A small percentage of the mature neurons expressed functional TRPV1, a cation channel involved in peripheral pain sensation that is activated by capsaicin. TRPV1 is expressed on a population of unmyelinated, slowly conducting C-fibers that express neuropeptides^34^, including substance P (TAC1) which is expressed during the differentiation into neural crest cells and remains expressed in the mature iPSC-SNs. The iPSC-SNs also have functional P2X3 channels, which are activated in response to ATP and lead to increased sensitivity to noxious stimuli^35^. The mature sensory neurons also exhibit a strong glutamate response, consistent with the expression of multiple metabotropic glutamate receptors in sensory neurons^36, 37^. Based on the expression of sensory neuron specific or enriched genes and the functional properties of the iPSC-SNs, they are considered a robust model for the study of peripheral neurotoxicity.

Microtubule targeting agents (MTAs) are the largest class of chemotherapeutic agents associated with sensory peripheral neuropathy. MTA-associated neuropathy demonstrates axonal degeneration that may be reversible in some individuals following removal of the agent, highlighting a mechanism distinct from other cytotoxic effects of MTAs^5, 38^. An *in vitro* model should reflect these features, including conditions where neurite retraction occurs with minimal cytotoxicity, as demonstrated in this study. The iPSC-SNs are mature neurons with complex neurite networks that can be stained for the neuron specific βIII-tubulin. Degeneration of the neurites showed dose- and time-dependent changes in response to paclitaxel exposure, consistent with the ‘dying back’ phenotype attributed to microtubule disruptors that leads to loss of re-innervation of the epidermis and CIPN symptoms^5^. A similar neurite retraction phenotype was quantified in an ESC-derived sensory neuron model of peripheral nerve injury^32^. In recent years, studies of chemotherapy neurotoxicity have been performed in commercially available neurons from iPSCs (e.g., iCell Neurons^®^ and Peri.4U Neurons^®^)^9–16, 19, 39^. In these cases, cells are typically tested shortly after plating and the outgrowth of neurites is measured using high content imaging. While neurite outgrowth is a common measurement of the neurodevelopmental effects of chemicals^40, 41^, the mature neurite networks investigated in this study more accurately reflect the conditions during exposure to paclitaxel in humans.

Optimal *in vitro* systems for studying chemotherapy neurotoxicity will be compatible with multiple measurements for assessing toxicity. The current results demonstrate that the iPSC-SNs are sensitive to the effects of paclitaxel on mitochondrial function and neuronal excitability. Paclitaxel-induced peripheral neuropathy has been associated with mitochondrial dysfunction and ATP deficits in both peripheral nerves and DRG neurons of paclitaxel-treated rats^42–44^. However, none of these paclitaxel-evoked changes could be replicated following *in vitro* paclitaxel exposure to naive DRG neurons^42^. In our iPSC-SN model, paclitaxel treatment significantly decreased the mitochondrial membrane potential, an indicator of mitochondrial dysfunction. Paclitaxel also disrupted axonal transport of mitochondria in a time-dependent manner, with long exposure to paclitaxel impairing both retrograde and anterograde movement. The transport of mitochondria along axons is essential for supplying ATP to sites of high energy demand, such as neurite ends, and mitochondria function is critical for intracellular Ca^2+^ homeostasis^45^. *In vivo* studies have demonstrated swollen and vacuolated mitochondria in axons of peripheral nerves that are linked to painful paclitaxel-induced neuropathy^46, 47^. While quantification of mitochondria transport was not possible in the current studies due to cell density requirements for long-term maintenance of function, measurement of mitochondrial membrane potential can be performed in a high throughput manner, providing opportunities to screen for reversal agents and to mechanistically investigate the pathways critical for the observed mitochondrial effects of chemotherapeutics.

Sensory neurons differentiated and induced from ESCs and iPSCs were positive for glutamate and characterized as excitatory glutamatergic neurons^26^. This is consistent with RNA-seq data showing strong expression of multiple ionotropic glutamate receptors and weak expression of limited metabotropic glutamate receptors in human iPSC-induced sensory neurons^24^ as well as detection of ionotropic glutamate receptors in peripheral axons of human skin^48^. Glutamate-induced excitability was detected in our iPSC-SNs and treatment with paclitaxel altered this excitability. Short-term exposure to paclitaxel increased glutamate excitability while longer-term paclitaxel exposure led to reduced excitability. The role of ionotropic and metabotropic glutamate receptors in paclitaxel-induced peripheral neuropathy is not well understood and most studies have focused on NMDA receptor activation in paclitaxel-induced pain^49, 50^. The current findings warrant further investigation to understand the time-dependent changes in response to paclitaxel and further expand the use of this iPSC-SN model for mechanistic studies of drug-induced changes in neuron excitability.

There are several advantages of the current iPSC-SN model to study chemotherapy-induced neurotoxicity compared to similar studies in iCell Neurons^®^ and Peri.4U neurons^®9–16, 19, 39^. The iPSC-SNs are derived from a well-characterized and widely used iPSC line^25^, with a known genetic background that can be edited for mechanistic investigation of specific genes and for the introduction of human polymorphisms that are associated with chemotherapy-induced peripheral neuropathy. Other iPSC-derived sensory neurons have proven useful for the study of peripheral nerve injury, mechanotransduction and neurodegeneration^30, 32, 51^. The differentiation and induction into sensory neurons are relatively robust processes and permit continuous culturing and generation of similarly acting iPSC-SNs over time. Methods described for cryopreservation of terminally differentiated sensory neurons can be applied in future large scale chemical and genetic screens^52^. The cells are also sensitive to drugs with varying mechanisms of action, including the microtubule stabilizers paclitaxel and docetaxel, the microtubule destabilizer vincristine and the proteasome inhibitor bortezomib, and will therefore be of value in dissecting the specific molecular mechanisms underlying drug-specific phenotypes. Finally, iPSC-derived sensory neurons can be cultured in the presence of Schwann cells^32, 53^ to investigate whether chemotherapeutics affect their interaction.

Although the iPSC-SNs offer a number of advantages over other *in vitro* models of chemotherapy-induced neurotoxicity, there remain limitations. While less expensive than commercially available iPSC-derived neurons, culturing and differentiation of iPSCs is still costly and requires long-term investment to maintain reproducible and readily available cultures for experimentation. Continuous culturing is resource intensive and requires a certain level of expertise. As discussed above, differences in seeding conditions and pluripotency status of the iPSCs prior to differentiation and minor changes in the differentiation and maturation protocol can significantly affect the composition of the derived sensory neurons. The use of mature sensory neuron populations has additional challenges related to the variability in extensive neurite networks that affect the ability to quantify morphology metrics with high content imaging.

In summary, the iPSC-SN model of chemotherapy-induced neurotoxicity described here is robust and displays multiple phenotypes associated with this dose-limiting toxicity. iPSC-SNs are an appropriate target cell to study CIPN and develop a pronounced axon degeneration in response to paclitaxel and other chemotherapeutics. This *in vitro* model of CIPN provides an attractive alternative to rodent models to elucidate relevant signaling pathways involved in chemotherapy-induced neurotoxicity. Future studies will use this platform for the investigation of genetic polymorphisms and gene pathways currently implicated in this toxicity^14, 16, 45, 54–56^. The iPSC-SNs will also be used for high throughput genetic and chemical screens to identify novel targets for the prevention or treatment of CIPN.

## Supporting information

Supplemental Movie, Vehicle

Supplemental Movie, Paclitaxel 0.1 uM

Supplemental Movie, Paclitaxel 1 uM

## Study Highlights

### What is the current knowledge on the topic?

Sensory peripheral neuropathy is a common and dose-limiting adverse event during chemotherapy. The lack of a molecular understanding of this toxicity limits options for its prevention and treatment.

### What question did this study address?

The current study tested whether sensory neurons differentiated from human induced pluripotent stem cells (iPSC-SNs) can be used to investigate chemotherapy-induced neurotoxicity, using paclitaxel as a model neurotoxic chemotherapeutic.

### What does this study add to our knowledge?

iPSC-SNs are a robust and reproducible model of paclitaxel-induced neurotoxicity. Treatment of iPSC-SNs with paclitaxel affects neurite networks, neuron excitability and mitochondrial function.

### How might this change clinical pharmacology or translational science?

This novel stem cell model of chemotherapy-induced neurotoxicity will be valuable for identifying genes and pathways critical for this toxicity and could be a useful platform for testing therapeutic approaches for treatment.

## Acknowledgements

The authors acknowledge helpful discussions with Dr. Sara Rashkin regarding the image analysis pipeline and with Dr. Bruce Conklin regarding stem cell models. They also acknowledge the valuable input on this manuscript from Drs. Josefina Priotti and Nura El-Haj.

## Author contributions

C.X., K.C.C. and D.L.K wrote the manuscript; C.X., K.C.C., T.B.S, F.F. and D.L.K designed the research; C.X., T.B.S, A.A.M, D.C, K.S and A.C.F performed the research; C.X., K.C.C. and D.L.K analyzed the data; K.C.C., J.K. and D.L.K contributed new analytical tools.

## Supplementary Materials

**Table S1. Compounds and reagents for differentiation and treatment**

**Table S2. Antibodies for immunocytochemistry**

**Table S3. Taqman probes for qPCR**

**Figure S1. Differentiation scheme for human iPSC-derived sensory neurons Figure S2. Quality control workflow for high content image analysis**

**Movie S1. Effect of paclitaxel on mitochondrial movement along the neurites**

