## Supplemental Tables for "Human Induced Pluripotent Stem Cell Derived Sensory Neurons are Sensitive to the Neurotoxic Effects of Paclitaxel"

**Table S1.** Compounds and reagents for differentiation and treatment

| **Compounds/Reagents** | **Supplier** | **Catalog Number** |
|---|---|---|
| DMEM/F12 basal medium | ThermoFisher | 11320033 |
| N2 Supplement | ThermoFisher | 17502048 |
| B27 Supplement | ThermoFisher | 12587-010 |
| Stemflex Medium Combo Kit | ThermoFisher | A3349401 |
| Poly-L-ornithine | Sigma-Aldrich | P3655 |
| Laminin mouse protein | Fisher Scientific | 23017015 |
| Febronectin | Fisher Scientific | 356008 |
| Matrigel Matrix | Corning | 356234 |
| Neuronal basal medium | ThermoFisher | 21103049 |
| Knockout DMEM | ThermoFisher | 10829018 |
| Knockout serum replacement | ThermoFisher | 10828028 |
| β-mercaptoethanol | ThermoFisher | 21985023 |
| Glutamax | ThermoFisher | 35050061 |
| MEM nonessential amino acids | ThermoFisher | 11140050 |
| LDN-193189 | Selleck Chemical | S7507 |
| SB431542 | Selleck Chemical | S1067 |
| CHIR99021 | Selleck Chemical | S1263 |
| SU5402 | Selleck Chemical | SU5402 |
| DAPT | Selleck Chemical | S2215 |
| Recombinant Human NT3 | PeproTech | 450-03 |
| Recombinant Human GDNF | PeproTech | 450-10 |
| Recombinant Human BDNF | PeproTech | 450-02 |
| Recombinant Murine beta-NGF | PeproTech | 450-34 |
| Y27632 | Selleck Chemical | S1049 |
| α,β-me-ATP | Sigma-Aldrich | M6517 |
| Capsaicin | EMD Millipore | 211275 |
| Glutamate | Sigma-Aldrich | G3291 |
| KCl | Sigma-Aldrich | P5405 |
| Mitomycin C | Sigma-Aldrich | M4287 |
| Paclitaxel | Sigma-Aldrich | T7402 |
| Docetaxel | Sigma-Aldrich | 01885 |
| Vincristine | Sigma-Aldrich | V8388 |
| Bortezomib | Selleck Chemicals | S1013 |
| Hydroxyurea | Sigma-Aldrich | H8627 |

**Table S2.** Antibodies for immunocytochemistry

| Antibody Name | Dilution | Host | Supplier | Catalog Number |
|---|---|---|---|---|
| SOX10 | 1: 500 | Goat | Santa Cruz | SC17342 |
| PAX6 | 1: 500 | Rabbit | Covance | PRB-278P |
| TUBB3 | 1: 1000 | Mouse | Covance | MMS-435P |
| TUBB3 | 1: 1000 | Rabbit | Covance | MRB-435P |
| Peripherin | 1: 500 | Goat | Santa Cruz | SC7604 |
| Brn3A | 1: 500 | Mouse | Millipore | Mab1585 |
| TRPV1 | 1: 200 | Guinea pig | ThermoFisher | PA1-29770 |
| TRKA | 1: 50 | Rabbit | Alomone Labs | ANT-018 |
| TRKB | 1: 50 | Rabbit | Alomone Labs | ANT-019 |
| TRKC | 1: 50 | Rabbit | Alomone Labs | ANT-020 |
| Goat anti-mouse Ig G | 1: 1000 | Goat | ThermoFisher | A11029 |
| Chicken anti-goat Ig G | 1: 1000 | Chicken | ThermoFisher | A21468 |
| Goat anti-guniea pig Ig G | 1: 1000 | Goat | ThermoFisher | A11076 |
| Goat anti-rabbit Ig G (Alexa | 1: 1000 | Goat | ThermoFisher | A11011 |

**Table S3.** Taqman probes for qPCR

| Gene | Probe ID |
|---|---|
| NANOG | Hs02387400_g1 |
| OCT4 | Hs00999632_g1 |
| SOX10 | Hs00366918_m1 |
| PAX6 | Hs01088114_m1 |
| TUBB3 | Hs00801390_s1 |
| TAC1 | Hs00243225_m1 |
| NTRK1 | Hs01021011_m1 |
