## Supplementary figures and images for "Human Induced Pluripotent Stem Cell Derived Sensory Neurons are Sensitive to the Neurotoxic Effects of Paclitaxel"

### Figure S1

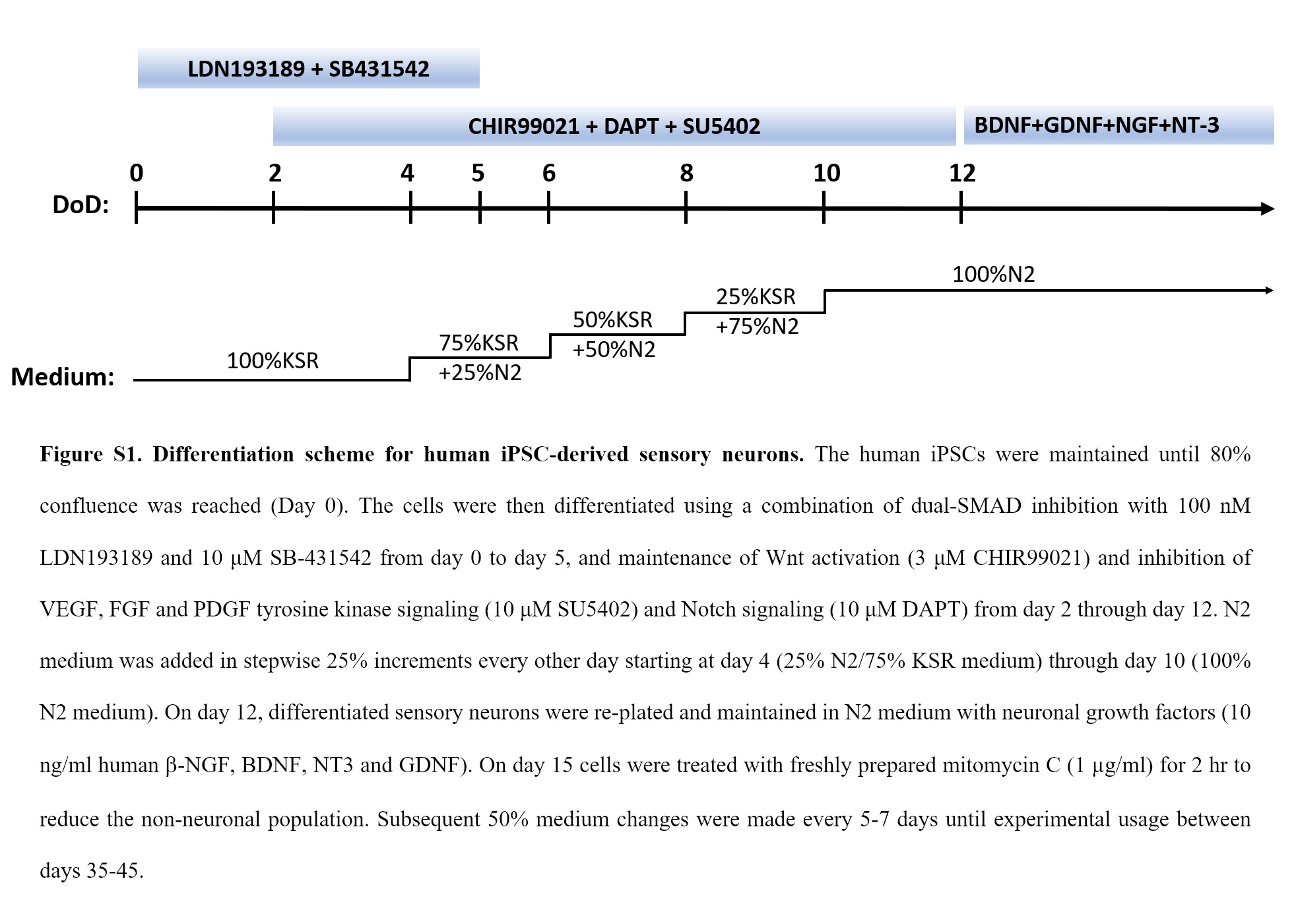

### Figure S2

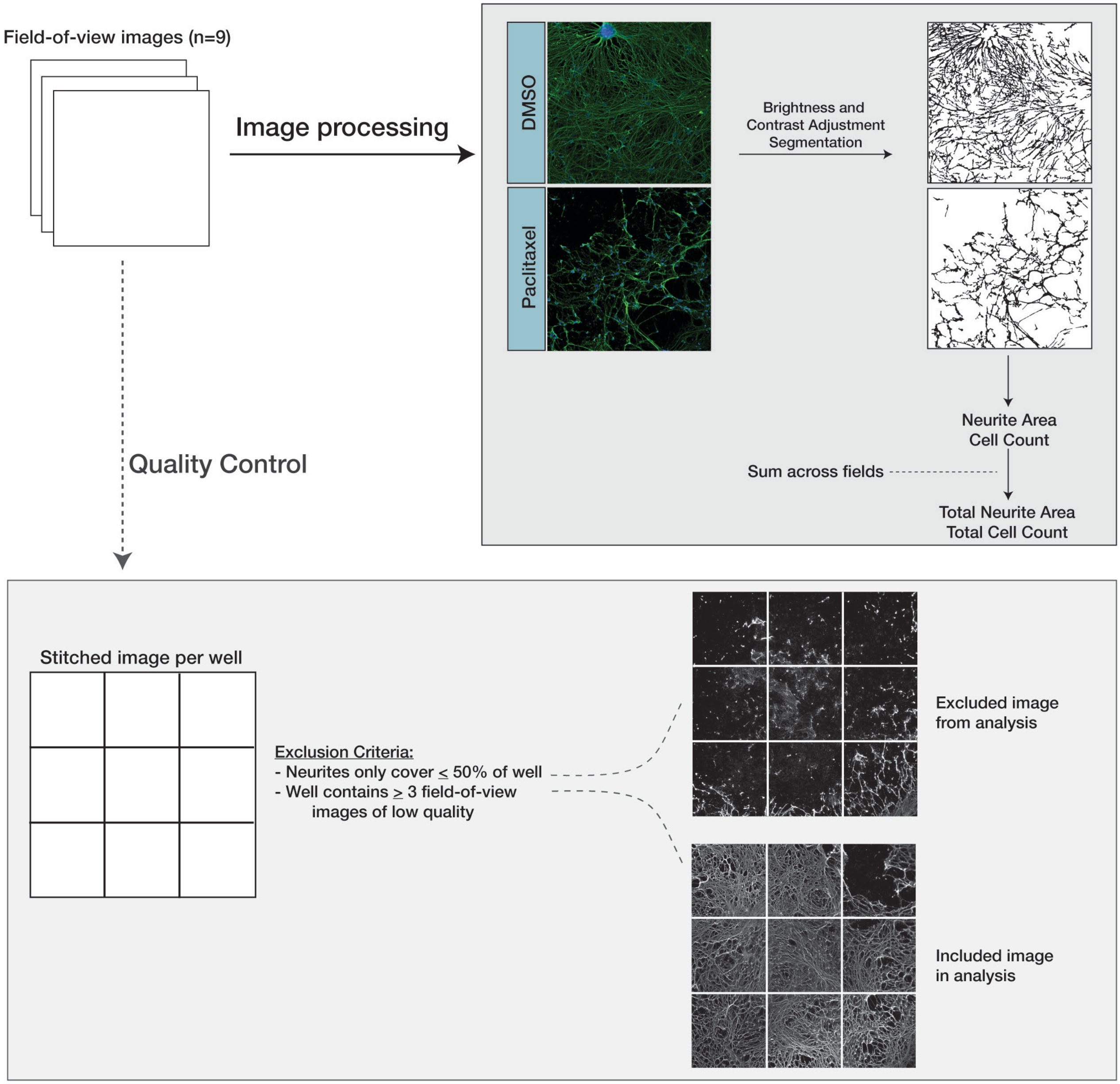


Figure S2. Quality control workflow of high content image analysis.
